# Mutations in S2 subunit of SARS-CoV-2 Omicron spike strongly influence its conformation, fusogenicity and neutralization sensitivity

**DOI:** 10.1101/2023.03.05.531143

**Authors:** Sahil Kumar, Rathina Delipan, Debajyoti Chakraborty, Kawkab Kanjo, Randhir Singh, Nittu Singh, Samreen Siddiqui, Akansha Tyagi, Sujeet Jha, Krishan G. Thakur, Rajesh Pandey, Raghavan Varadarajan, Rajesh P. Ringe

## Abstract

SARS-CoV-2 has remarkable ability to respond to and evolve against the selection pressure by host immunity exemplified by emergence of Omicron lineage. Here, we characterized the functional significance of mutations in Omicron spike. By systematic transfer of mutations in WT spike we assessed neutralization sensitivity, fusogenicity, and TMPRSS2-dependence for entry. The data revealed that the mutations in both S1 and S2 complement to make Omicron highly resistant. Strikingly, the mutations in Omicron S2 modulated the neutralization sensitivity to NTD- and RBD-antibodies, but not to S2 specific neutralizing antibodies, suggesting that the mutations in S2 were primarily acquired to gain resistance to S1-antibodies. Although all six mutations in S2 appeared to act in concert, D796Y showed greatest impact on neutralization sensitivity and rendered WT virus >100-fold resistant to S309, COVA2-17, and 4A8. S2 mutations greatly reduced the antigenicity for NAbs due to reduced exposure of epitopes. In terms of the entry pathway, S1 or S2 mutations only partially altered the entry phenotype of WT and required both sets of mutations for complete switch to endosomal route and loss of syncytia formation. In particular, N856K and L981F in Omicron reduced fusion capacity and explain why subsequent Omicron variants lost them to regain fusogenicity.

## Introduction

The best response to the COVID-19 pandemic has been the development of vaccines which helped fight the pandemic and reclaim normalcy. However, the emergence of new and highly transmissible variants has jeopardized the vaccine efficacy (1–7). Several SARS-CoV-2 variants categorized as variants of concern (VOC) and variants of interest (VOI) have emerged over the past year that influenced the trajectory of the pandemic(2, 8–10). The latest variant B.1.1.529 (Omicron) and its newer sub-lineages (BA.2, BA.4, BA.5, BQ.1, and XBB.1) have acquired resistance to vaccine-induced immunity and has been infecting large population worldwide (11–15). It is highly plausible that there is still much more genetic space that SARS-CoV-2 can explore to navigate through host immunity without compromising transmissibility(16–19).

The spike protein is composed of S1 and S2 subunits in trimer-of-heterodimers meta-stable state. The protein is conformationally flexible and undergoes large structural transitions after receptor binding leading to membrane fusion and virus entry(20, 21). The S1 is surface protein and harbors key functional domains which are well-exposed to the immune system(22–28). The antibodies are frequently directed against receptor binding domain (RBD) and N-terminal domain (NTD) which antagonize the binding of spike to cellular receptor. These virus neutralizing antibodies (NAbs) exert selection pressure on these domains giving rise to progeny with escape mutations(17, 29–31). The most resistant VOC prior to the emergence of Omicron was Beta which acquired resistance to convalescent or vaccine sera through only 10 mutations; amongst which E484K is most important and can alone drive much of neutralization resistance(32–34). Omicron, however, acquired 28 mutations in S1 including 15 in RBD, and 6 mutations in S2. While the S2 subunit is conformationally stable in the pre-fusion state of spike, the S1 subunit is inherently dynamic(21, 35, 36). Three RBDs on each trimer can transition from closed “down” to open “up” conformation and determine the sensitivity to neutralizing antibodies and enable binding to cellular receptor. Similarly, NTD, although heavily protected by glycans, has epitopes for neutralizing antibodies and its conformational dynamics influence their binding to the “supersite”(26, 37, 38). The open and closed transitions of these domains create intrinsic heterogeneity for the virus to exploit against neutralization. The receptor binding motif (RBM) which is occluded in RBD’s “down” conformation is exposed in the “up” conformation and favors interaction with ACE-2 (21, 35, 36, 39). Binding to the receptor follows proteolytic cleavage of the site in S2 and triggers massive conformational transition and insertion of fusion peptide (FP) into the cellular membrane that leads to fusion of the virus and cell membranes(40, 41). Although the S1 is flexible and dynamic the RBD-RBD, NTD-RBD interactions are tightly regulated by direct contacts or through the structural linkers (36, 42, 43). The NAb resistance mutations are selected based on the principle that they do not affect the interaction with receptor and fusion with the host cell but prevent binding of NAbs elicited by previous infections (44–46). Omicron has acquired several mutations in NTD and RBD as well as SD-1 and SD-2. In addition to that, and unlike previously known VOCs, Omicron has acquired 6 mutations in S2. These mutations alter the conformational states of S1 domains and also the S2 subunit which in turn impact antibody binding to their cognate epitopes(47).

In the present work, we have studied the effects of mutations in Omicron spike on the virus’s ability to resist neutralizing antibodies, and on the entry pathway. The mutations in S2 subunit have persisted in all the omicron variants suggesting that they have a strong functional role. The immune pressure is less on S2 as the antibodies against S2 subunit are induced in much lower titer and generally with less potency relative to S1 directed antibodies. Therefore, the selection of mutations in S2 must have indirect overall effects on the conformation and function of spike protein. Identification of the mutations and understanding the mechanism of NAb resistance is critical for vaccine design. Similarly, the pathway of virus entry has implications for pathogenicity. Omicron is less pathogenic than earlier VOCs as it doesn’t fuse at the plasma membrane (PM) and does not form syncytia, unlike Delta which is highly fusogenic and much more pathogenic(48, 49). While Omicron is poorly replicating due to its altered entry pathway, the new sub-lineages of Omicron tend to reverse this phenotype by acquiring novel mutations (50). Therefore, understanding the mechanism and relationship between neutralization resistance and virus entry of Omicron is important and can provide insights into SARS-CoV-2 pathogenicity and vaccine design.

## Results

### Omicron is resistant to vaccine sera but breakthrough infection boosts neutralization titers

In India, two vaccines were approved in 2020 in the national program of vaccination – Covishield and Covaxin, and most of the population is vaccinated using these vaccines. Covishield is a chimpanzee adenovirus expressing the SARS-CoV-2 spike and it is produced by Serum Institute of India. The Covaxin is a whole virus inactivated vaccine developed by Bharat Biotech, India(51, 52). The sera from vaccinees were collected after two doses of the same vaccine between 3-5 weeks post vaccination and used for the assessment of neutralization titers. The Covishield vaccine sera (n = 33) neutralized pseudovirus expressing WT spike potently (GMT-857) compared to other VOCs (Figure 1A). The neutralization titers significantly decreased for Alpha (P=0.003), Beta (P<0.0001), Delta (P<0.0001), and Omicron (P<0.0001). Omicron was the most resistant (GMT 81) followed by beta (GMT 136) and only 13 and 17 out of 33 sera neutralized these variants respectively with serum neutralization titer of >50. Thus, as reported in several other studies E484K present in Beta probably drove resistance to serum Nabs whereas E484K and several other mutations throughout the spike of Omicron afforded the greatest resistance to neutralization. The Covaxin sera (n=10) were weakly neutralizing compared to Covishield sera (Figure 1A, B). The GMT of neutralization of Covaxin sera against the WT and Alpha variant was 208 and 211 respectively whereas it was significantly reduced for Beta and Delta (Figure 1B). None of the sera neutralized Omicron but three sera neutralized the Beta variant at serum dilution of >100.

**Figure 1:**
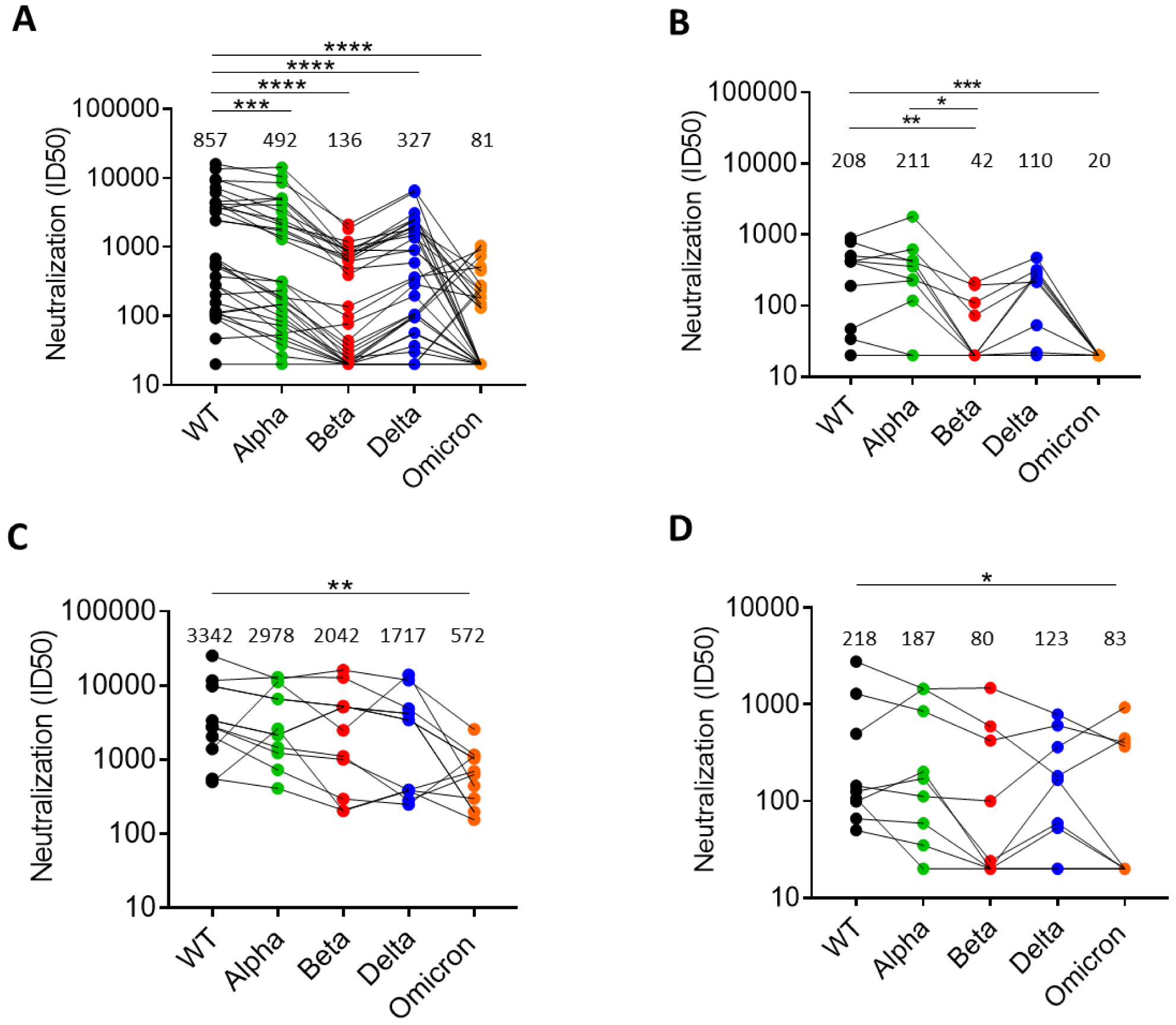
Neutralizing antibody response induced by vaccines to SARS-CoV-2 variants: The NAb response assessed in pseudovirus neutralization assay against five SARS-CoV-2 variants of concern induced by A) Covishield sera (n=33) B) Covaxin sera (n=10) C) Covishield followed by breakthrough infection sera (n=8) D) Covaxin followed by breakthrough infection sera (n=9). The geometric mean titer is shown for each pseudovirus above each column. The statistical significance of differences between the groups was calculated by using Wilcoxon matched-pairs signed rank test and two-tailed p value are indicated as **** (P<0.0001), *** (P<0.003

We also collected sera samples from vaccinated people who became infected with SARS-CoV-2 after >3 months post-vaccination. These individuals with hybrid immunity (HI) were different from only-vaccine group, therefore, comparative analysis was cross-sectional. The sera were collected during the third wave in India in which Omicron was the dominant variant circulating and Delta being the subdominant variant. The median neutralization titers in Covishield vaccinees (n=12) were increased against all the variants tested (Figure 1C). All the sera also neutralized Omicron variant with moderate titers although the titer was significantly lower than WT virus (P=0.008) (Figure 1C). The titers significantly increased in HI group compared to vaccine group against Alpha, Beta, Delta, and Omicron (Figure S1). The neutralization titers in Covaxin vaccinees (n=10) were also increased against all the variants tested, although, 5 sera still could not neutralize beta and Omicron variants (Figure 1D). Overall, the titers were much lower in Covaxin than Covishield, but statistical significance could not be obtained due to limited number of samples.

### The neutralization resistance of Omicron is contributed by both S1 and S2 subunit mutations

There are 34 mutations in Omicron spike out of which 28 are present in S1 and 6 are in S2 subunit. To investigate the contribution of these mutations to neutralization resistance of Omicron we made the chimeric spikes by replacing S1 or S2 subunits of WT with that of Omicron (Figure 2A). In the third construct, two Omicron substitutions N679K and P681H near furin cleavage site (FCS) were also made in WT spike background (Figure 2A). We assessed the neutralization sensitivity of WT and chimeric spike pseudoviruses bearing WT-Omi-S1, WT-Omi S2, and WT-Omi-FCS to selected potent Covishield sera (Figure 2B). The pseudoviruses bearing WT or WT-Omi-FCS spike were similarly sensitive to these sera but the WT-Omi-S1 pseudovirus was resistant and its degree of resistance was equivalent to Omicron pseudovirus (Figure 2B & C). Whereas, WT-Omi S2 pseudovirus was sensitive to all sera but its sensitivity was significantly reduced when compared with WT (P= 0.012) (Figure 2C). These results suggested that most of the neutralization resistance was driven by the mutations in S1 domain and that is probably because they directly disrupt the sequence of epitopes or their conformations. The mutations in S2, albeit modestly, also contributed to resistance. The contribution of S2 mutations in neutralization resistance was surprising as S2 is poorly immunogenic and there is not expected to be selection pressure on this subunit.

**Figure 2:**
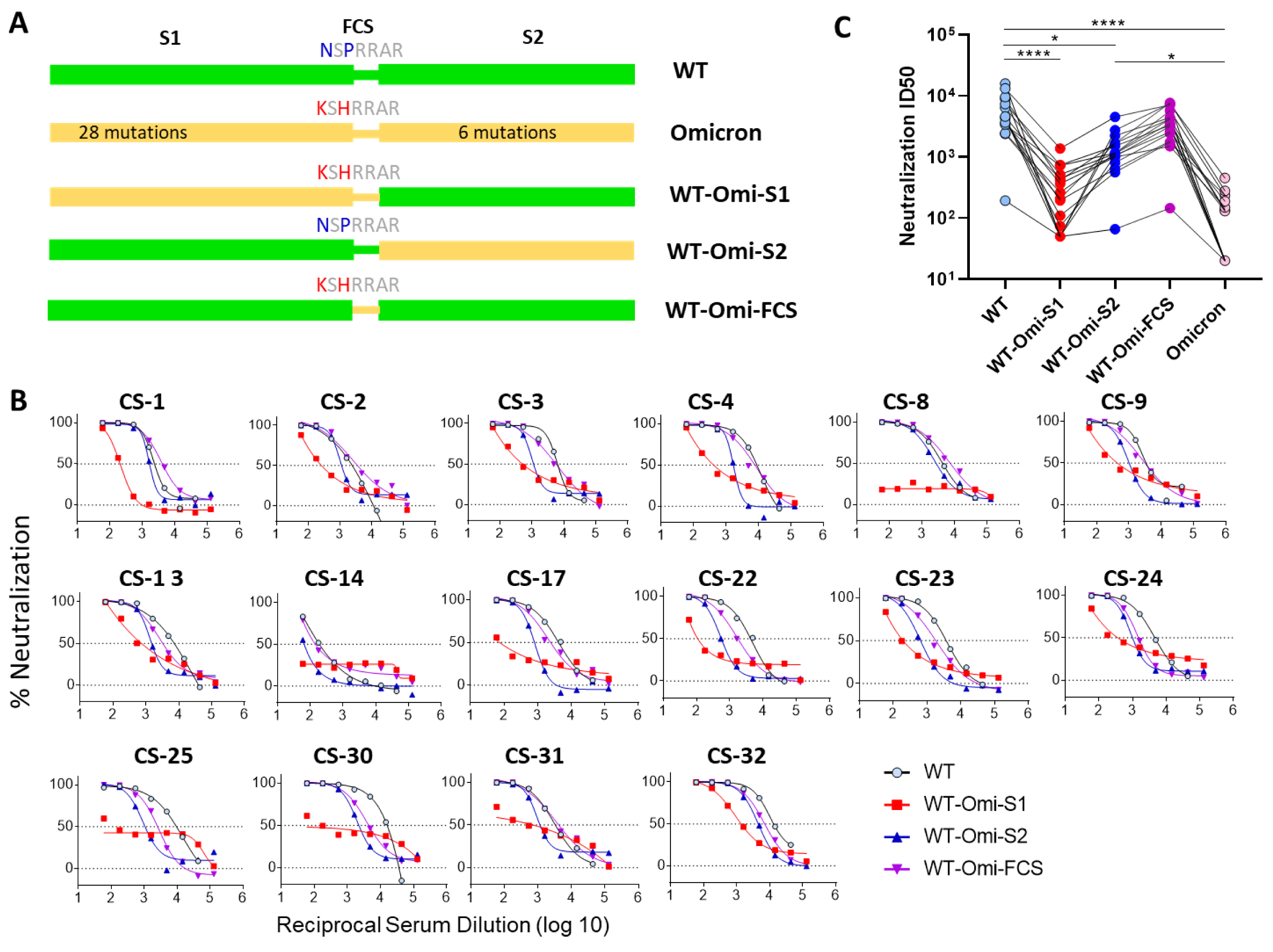
Neutralization sensitivity of WT and mutant spike-expressing pseudoviruses to Covishield vaccine sera: The Omicron mutations in S1, S2 subunit, and two mutations near furin cleavage site (N679K, P681H) were separately transferred in WT spike A) schematic of the spike protein shown for each parental and mutant spike protein B) Neutralization of pseudoviruses representing WT or mutant spike by select high-titer Covishield vaccine sera and percent neutralization plotted against serially diluted sera C) Inhibitory dilution-50 (ID50) for the same sera in B plotted against parental and mutant viruses. The statistical significance of difference in neutralization titers was calculated as mentioned in Figure legend 1.

We considered two possibilities that might have driven the neutralization resistance via S2 mutations (i) there are low-titer antibodies in the sera targeting the S2 subunit and the mutations in S2 affected their binding to epitopes in that subunit. (ii) the mutations in S2 subunit indirectly altered the presentation of NAb epitopes in S1. To examine the second possibility, we compared the neutralization sensitivity of WT and WT-Omi S2 pseudoviruses, both of which carry S1 of WT spike, to mouse sera raised by RBD immunogens(53, 54). These RBD immunogens were based on WT-spike sequence with or without stabilizing mutations (53, 54). The mice sera potently neutralized both WT and WT-Omi S2 pseudoviruses (Figure 3A). However, the neutralization potency against WT-Omi S2 was reduced for most of the sera (Figure 3A & B). The geometric mean titers for the WT and WT-Omi S2 virus were 9982 and 6847 respectively (Figure 3B). Taken together, the ID50 values were significantly reduced for WT-Omi S2 compared to WT pseudovirus (p <0.0001). To further consolidate the role of S2 mutations, we analysed the neutralization of pseudovirus bearing Omicron or WT-Omi-S1 spike, the latter with S2 mutations reverted to WT (Figure 2A). We used sera from mice which were immunized with Omicron-specific RBD antigen (ie: RBD contained Omicron mutations). These sera potently neutralized both Omicron and WT-Omi-S1, however, the latter was significantly more sensitive than the former (P=0.03) (Figure 3C). Taken together, these data suggested that the mutations in S2 subunit modulated the neutralization sensitivity to RBD antibodies.

**Figure 3:**
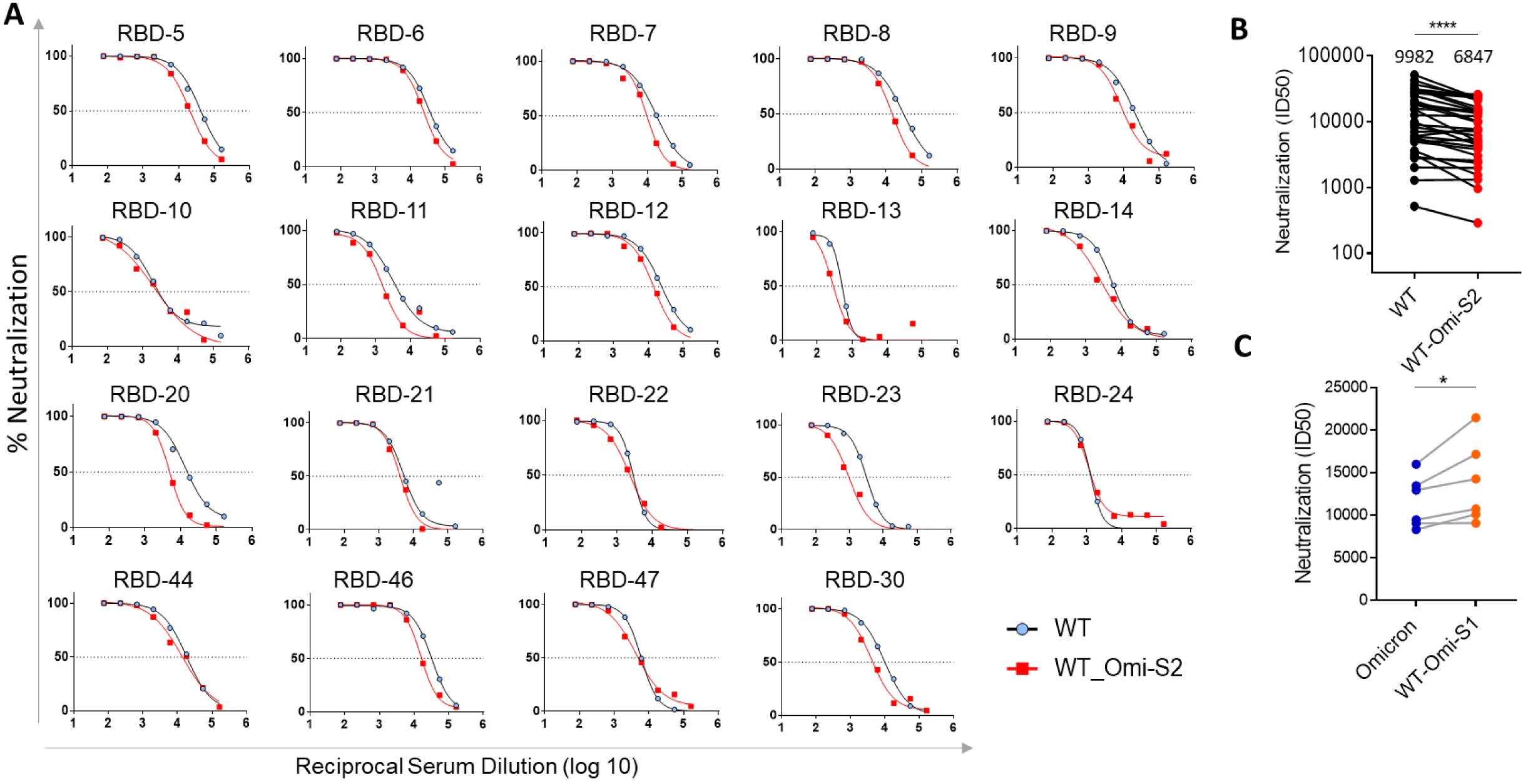
Neutralization sensitivity of WT and WT-Omi S2 to RBD sera: A) The neutralization curves are shown for WT and WT-Omi S2 which is a mutant containing omicron-S2 mutations in the background of WT spike. B) The neutralization titers against both the viruses are shown for mouse sera (n=34) which were raised against WT-RBD immunogens and sera were drawn after two doses of RBD vaccine. The GMT titer is shown above each column. C) The neutralization titers against WT and WT-Omi S2 pseudoviruses are shown for mouse sera (n=06) raised against Omicron-RBD immunogen

### The mutations in S2 affect the neutralization by monoclonal antibodies specific to NTD and RBD

Next, we wanted to see the effect of mutations in S2 subunit on neutralization by antibodies that target NTD and distinct epitopes in RBD. We used a panel of pseudoviruses consisting of parental (WT and Omicron), chimeric (WT-Omi-S1, WT-Omi S2), FCS mutant (WT-Omi-FCS), and WT virus with individual S2 mutations (Figure 4). We used COVA2-17, 4A8 (NTD), and COVA2-15, S309 (RBD) antibodies. In addition, we also used a pooled serum from mice, which were immunized with RBD immunogens, and recognize multiple epitopes in RBD (54). The neutralization sensitivity of WT-Omi S2 was unchanged for COVA2-15 compared to WT. However, the neutralization sensitivity to S309 was reduced for WT-Omi S2 and some S2 mutants, particularly, WT-D796Y and WT-L981F markedly reduced the sensitivity to this antibody (Figure 4A, C). In WT spike background, the D796Y alone enhanced neutralization resistance significantly Covishield or convalescent human sera (Figure S2A, B). The reversion of this mutation in Omicron spike significantly enhanced the neutralization by RBD sera reinforcing the role of this mutation in neutralization (Figure S2C). For the polyclonal serum pool, WT-Omi S2 and some of the point mutations also became slightly resistant (Figure 4B). The NTD antibodies 4A8 and COVA2-17 neutralized WT virus potently, although the maximum neutralization achieved was in the range of 70-80%. These antibodies did not neutralize Omicron and WT-Omi-S1 due to sequence change in epitopes and directly affecting their binding. Interestingly, WT-Omi S2 became completely resistant to COVA2-17 and partially resistant to 4A8 (Figure 4A). The single S2 mutations also variably altered the neutralization pattern for NTD antibodies - several mutations reduced maximum neutralization to <50% (Figure 4C). However, the neutralization by S2-specific antibodies (CC40.8 and CV3.25) was not changed (Figure 4A). This data revealed that the mutations in S2 indirectly modulated the exposure of epitopes in NTD as well as RBD thereby affecting neutralization.

**Figure 4:**
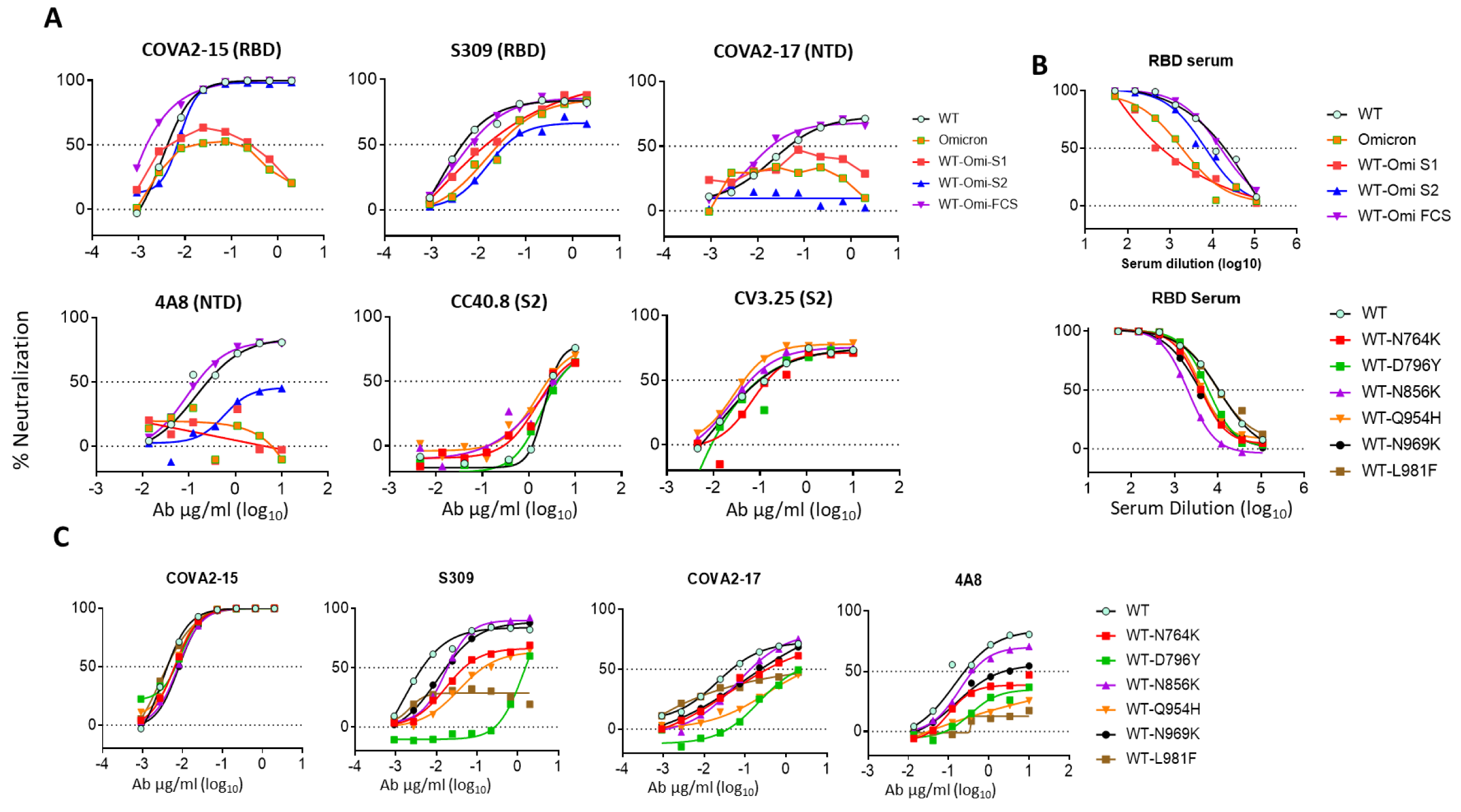
Neutralization resistance to monoclonal neutralizing antibodies conferred by Omicron mutations: A) The neutralization pattern of parental or mutant pseudoviruses to RBD or NTD or S2-specific monoclonal antibodies B) The neutralization pattern of pseudoviruses in A and WT mutants (individual mutations) by RBD-specific serum pool from vaccinated mice. C) Neutralization sensitivity of WT spike containing single S2 mutations to monoclonal antibodies

### The S2 mutations in omicron alter the exposure of epitopes in S1 subunit

The Omicron spike is relatively more stable and has a more compact configuration than WT or Delta spike (47, 55, 56). In addition to epitope disruptions by mutations in RBD and elsewhere in the S1 subunit, the slower transition from “down” to “up” conformation of RBD could be another reason why this variant is more resistant to RBD antibodies(47). The WT-Omi S2 spike showed reduced sensitivity to monoclonal antibodies targeting RBD and NTD, and also to the polyclonal sera raised by RBD immunogen. This result indicated that the mutations in S2 subunit modulate conformational flexibility and influence the exposure of epitopes in S1. To understand the mechanism of reduced exposure, we assessed the kinetics of neutralization. The virus was incubated with antibody for various time period before adding to the target cells. S309 neutralized WT virus efficiently in 10-minute incubation period and neutralization increased slightly at 30- and 120-minutes (Figure 5A). In contrast, WT-Omi S2 was neutralized slowly but neutralization substantially increased at longer incubation times (Figure 5A). The reduction in S309 IC_50_ values from 10-minute to 120-minute incubation was by 4-fold and 32.7-fold for WT and WT-Omi S2 respectively. Similarly, 4A8 IC_50_ for WT and WT-Omi S2 was reduced by 4.34- and >43-fold respectively from 10-minute to 120-minute incubation (Figure 5B). However, for COVA2-15, which is a cluster-I antibody targeting RBM, the neutralization increased moderately and only 3- and 6.33-fold reduction in IC_50_ was seen for WT and WT-Omi S2 respectively. (Figure 5A & B). Conversely, the same experiment was done with Omicron and BA.5 by comparing with their mutants (reversion of S2 mutations to WT) WT-Omi S1 and WT-BA.5 S1 respectively. S309 neutralized WT-Omi-S1 and WT-BA.5 S1 faster than their counterparts suggesting that the S2 mutations influenced the exposure of epitopes in RBD (Figure S3). Next, the overall effect of reduced exposure on neutralization sensitivity was assessed using various sera samples. The human vaccine sera neutralized WT-Omi S2 less well than WT (Figure 5C). Conversely, Omicron BA.1 or BA.5 with reverted S2 mutations (WT-Omi S1 or WT-BA.5 S1) were better neutralized than their parental pseudoviruses by the Omicron-RBD-specific sera (Figure 5D, E). As to how the S2 mutations might be influencing the neutralization, we further analyzed the antigenicity of WT and WT-Omi S2 spikes by FACS assay or ELISA using soluble spike protein. The S309 and COVA2-15 both bound better to WT than WT-Omi S2 tracking with the neutralization results (Figure 5F, G).

**Figure 5:**
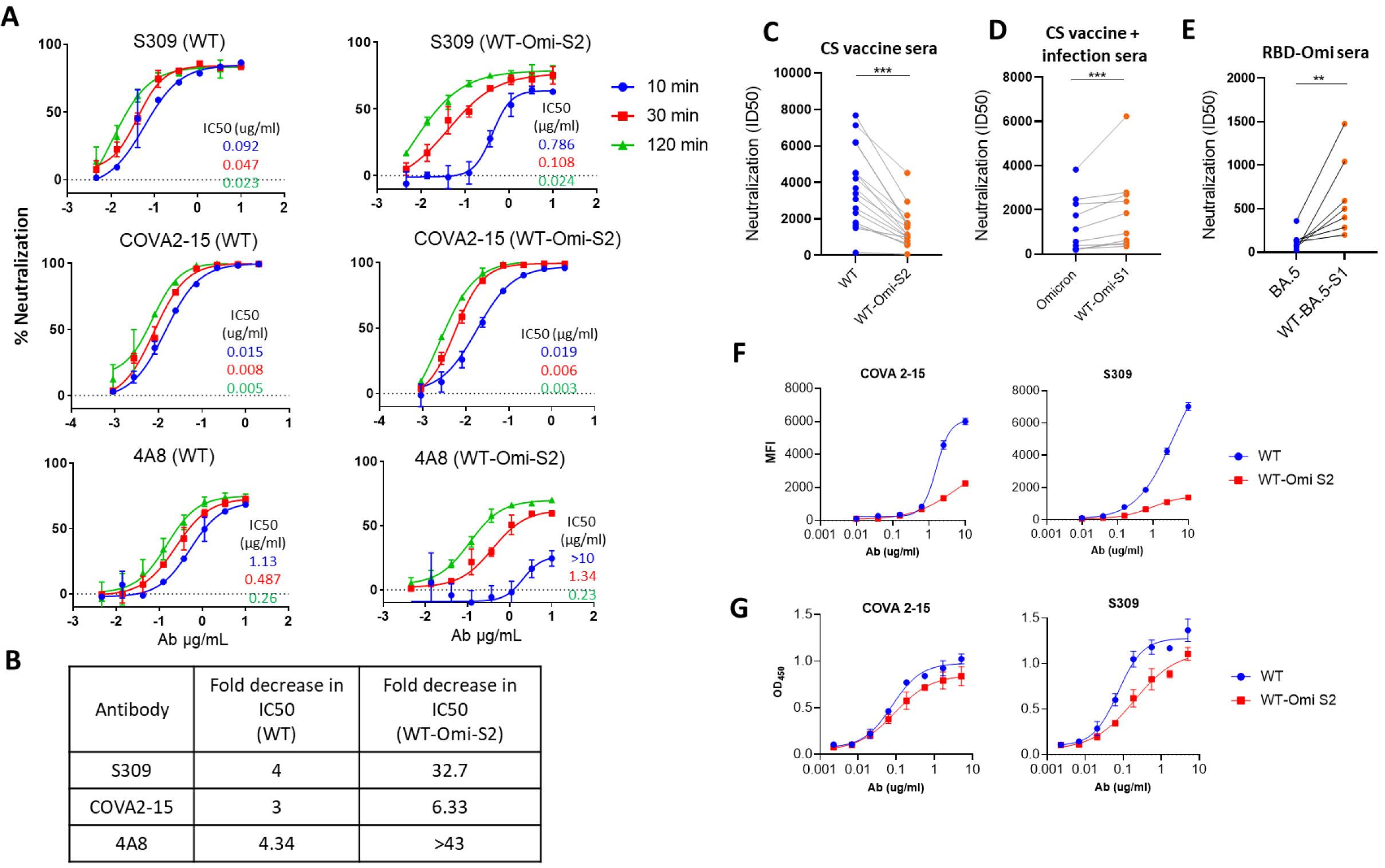
Neutralization kinetics of WT and WT-Omi S2 pseudoviruses by monoclonal NAbs: A. Neutralization curves are shown for WT and WT-Omi S2 pseudoviruses by Nabs incubated with virus for 10 or 30 or 120 minutes. The IC50 values (ug/ml) are shown in inset derived from respective neutralization curves and indicated by same color as used for respective neutralization curves. B. The IC50 values obtained at 10-minute and 120-minute incubation time were calculated. The fold decrease in IC50 value from 10 to 120-minute incubation for indicated antibodies is shown. C. Comparison of sensitivity of WT and WT-Omi S2 pseudovirus to Covishield vaccine sera. D. Sensitivity of Omicron BA.1 and WT-Omi S1 pseudovirus to Covishield vaccine followed by breakthrough infection sera. E. Neutralization sensitivity of Omicron BA5 and WT-BA5 S1 to mice sera elicited by Omicron-RBD immunogen. F, G. The binding of RBD NAbs to WT and WT-Omi S2 spikes expressed on 293T cells or as soluble ectodomain with K986P, V987P (2P) stabilizing mutations. Binding is shown as mean fluorescence intensity (MFI) for FACS assay and OD at 450nm for ELISA assay.

### The route of viral entry and fusogenicity is determined by sequence changes in S1 and S2

A major adaptation in Omicron compared to earlier VOCs is that it is less dependent on TMPRSS2 for the cleavage of S2’ site (second cleavage site in S2) and release of fusion peptide. Due to the non-requirement of TMPRSS2-mediated cleavage of Omicron spike at the cell surface, it is not primed at the cell surface and virus does not fuse at this location(49). The S2’ site is conserved across the variants and we hypothesized that the mutations in S2 subunit of Omicron might affect the cleavage by TMPRSS2 and thereby the fusion process at PM. We used the same panel of pseudoviruses as described in Figure 2 to study the effect of S2 mutations on the entry-related features. (Figure 6).

**Figure 6:**
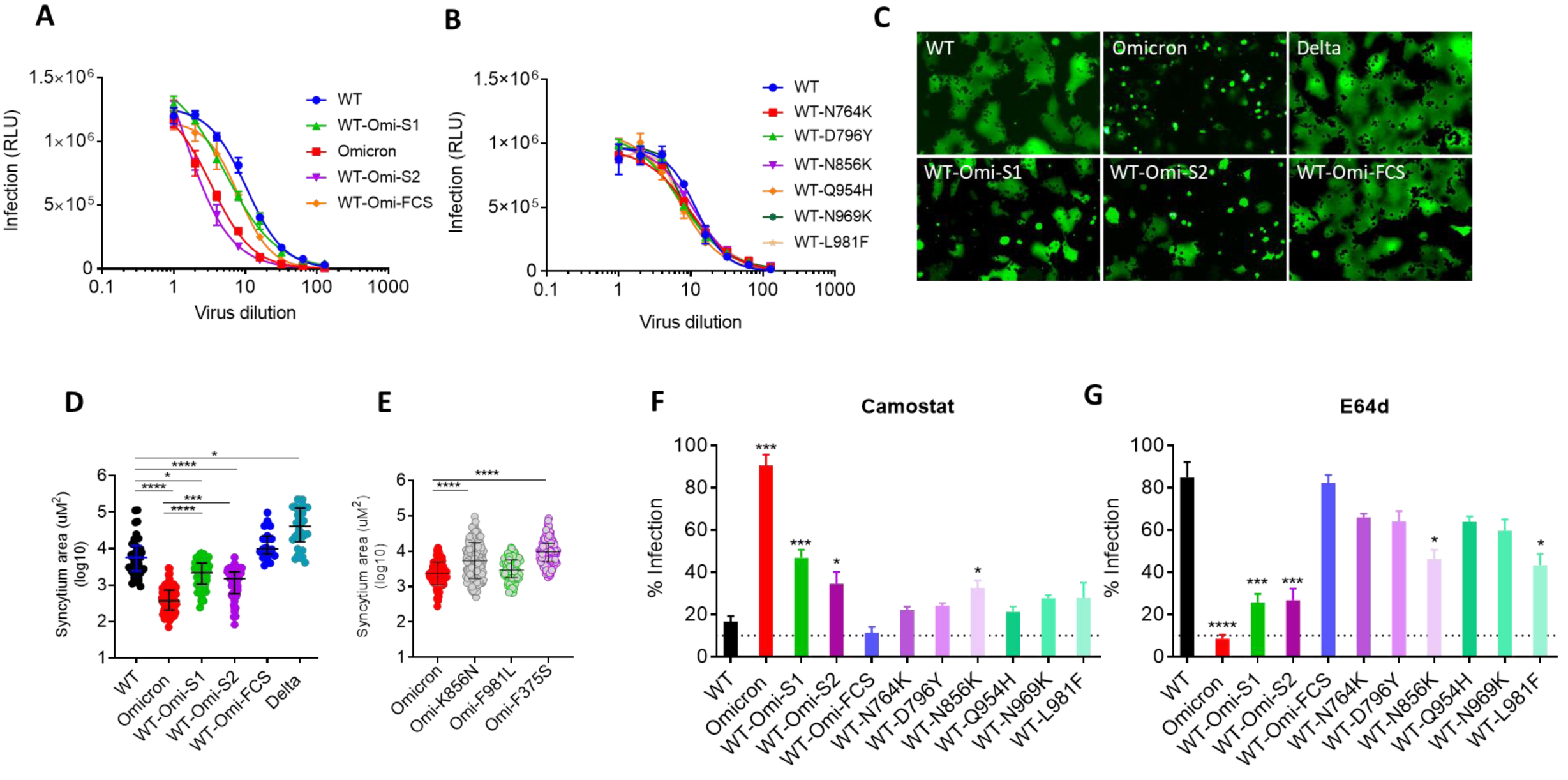
Entry of pseudoviruses bearing various spike proteins: Freshly prepared pseudoviruses were used for the infection of 293T-hACE2-TMPRSS2 cells by serially diluting the virus. A, B. Infectivity (relative luminescence unit -RLU) is shown against various dilutions of pseudovirus expressing parental and mutant spikes. C) 293T cells expressing Spike and GFP were added onto the monolayer of Vero-TMPRSS2. Shown are the images of syncytia formed by Spike variants. D, E. The size of the syncytia measured in uM^2^ (micrometer squared) plotted for indicated Spike variants. The statistical significance calculated using Kruskal-Wallis test and multiple comparison was done by Dunn’s multiple comparison test. F, G. Infectivity of pseudoviruses in the presence of TMPRSS2-inhibitor camostat or cathepsin-inhibitor E64D. The statistical analysis was done by using Kruskal-Wallis test and p values were calculated by Dunn’s multiple comparison post-test. All shown statistical significance values were derived in comparison with WT.

The infectivity of WT-Omi-S1 and WT-Omi-FCS mutant (N679K, P681H) was slightly reduced compared to WT pseudovirus but the infectivity of WT-Omi S2 was markedly reduced and was comparable to Omicron (Figure 6A). However, individual S2 mutations in WT background did not affect the infectivity suggesting that the set of six mutations in S2 collectively affect the infectivity (Figure 6B). The fusogenicity was highest for Delta followed by WT, and the Omicron variant showed least fusogenicity and displayed very few syncytia which were significantly smaller in size than WT (Figure 6C). The chimeric spike pseudoviruses WT-Omi-S1 and WT-Omi S2 showed reduced ability to fuse with receptor-expressing cells and size of their syncytia also was significantly reduced (Figure 6C, D). The fusogenicity of WT-Omi-FCS mutant was slightly enhanced and was comparable to Delta. The fusogenicity of WT containing individual Omicron S2 mutations showed similar fusogenicity as WT (Figure S4A). Conversely, reversion of S2 mutations in Omicron background was also evaluated. Here, the K856N and F981L clearly increased the fusogenicity of Omicron spike (Figure 6E, S4B). The F375S change which was previously reported by Kimura et al. for increased fusogenicity of Omicron was also confirmed for increased fusogenicity in this comparison (Figure 6E, S4B). Additionally, to assess the general effects of S2 mutations, Omicron BA.5 variant was compared with its mutant counterpart in which all four S2 mutations were changed to WT. The fusogenicity of mutant spike (WT-BA5-S1) was significantly increased (Figure S2D). Collectively, the data revealed that S2 mutations reduced the fusogenicity of Omicron lineage spikes.

We next assessed the effect of mutations on the virus entry mechanism. We used protease inhibitors camostat and E64d as inhibitors of TMPRSS2 (route-i) and endosomal cathepsins (route-ii) respectively to assess the dependence of these proteases on entry of SARS-CoV-2 variants. The infectivity of WT virus was reduced to baseline by camostat but was not affected by E64d. Conversely, the infectivity of Omicron was sensitive to E64d but not to camostat which suggested that WT mainly used TMPRSS2 and fused at PM whereas Omicron mainly used cellular cathepsins in endosomal compartment. WT-Omi-S1 and WT-Omi S2 both were partially sensitive to E64d and camostat. Compared to WT, the infectivity of WT-Omi-S1 and WT-Omi S2 was significantly higher in the presence of camostat or significantly lower in the presence of E64d (Figure 6F, G). The sensitivity of WT-Omi-FCS to both the inhibitors was similar to WT. The infectivity of all six single S2 mutants in the presence of camostat was slightly higher than WT but only for N856K was it significantly higher than WT. The infectivity of N856K and L981F was significantly reduced by E64d treatment which suggested that these two mutations enhanced endosomal entry of Omicron BA.1 more than other mutations in S2 (Figure 6F, G). The enhanced endocytosis of WT by these two mutations also corroborated the enhanced fusogenicity of reverse mutations in Omicron spike (i.e., the opposite effect) (Figure S4B). Overall, the mutations in either of the subunit only partially switch the entry pathway from PM to endocytosis but the mutations in both subunits complement for complete switch to endosomal route of entry.

## Discussion

In the present study, we investigated the role of mutations in S1 and S2 subunits of omicron spike protein. Omicron is by far the most mutated variant of SARS-CoV-2 since the onset of COVID-19 pandemic, and apart from acquiring neutralization resistance it also employs a different entry pathway. As there are several mutations in omicron spike it is unlikely that any single mutation or a smaller set of mutations would contribute to these phenotypes, instead it is a cumulative effect of several mutations(57). As expected, and because neutralization targets lie in the S1 subunit, the mutations in S1 drove most of the resistance due to epitope disruptions, however, mutations in S2 also afforded substantial resistance to neutralization. The S2 mutations are mostly buried in the spike structure, and are not part of prominent epitopes in S2, and did not alter the neutralization by S2-specific NAbs (58, 59). Therefore, resistance imparted by S2 mutations must be due to their allosteric effects on S1 epitopes. The set of six S2 mutations in WT spike reduced its neutralization sensitivity to monoclonal or polyclonal antibodies targeting the RBD and also resulted in marked resistance to S309, 4A8 and COVA2-17. The individual S2 mutations in the WT background also variably reduced the sensitivity to RBD and NTD antibodies, particularly, D796Y. This mutation is located close to the fusion peptide (FP) and may alter the FP dynamics and change the energetics of RBD position for reduced accessibility. This appears to be an additional strategy employed by viruses to gain neutralization resistance to antibodies targeting receptor binding domains (60, 61) Kemp et al. showed that D796H/D796Y frequently surfaced in viremia in an immunocompromised COVID-19 patient during plasma therapy(29). Schmidt et al. also revealed that SARS-CoV-2 acquired mutations, P792H and N801D, close to this region when virus was passaged in the presence of neutralizing sera underscoring the importance of this domain in regulating spike conformation (30).

Omicron spike protein represents a highly stabilized conformation due to improved contacts at the interface of S1 and S2 and at the interprotomeric S1-S1 and S2-S2 contacts which result in a compact configuration of omicron spike (55). The resistance to NAbs can be partially brought about by mutations in RBD and NTD whereas mutations in S2 core are used as leverage to compact the overall configuration of the spike to reduce the exposure of epitopes in RBD and NTD. (47). Reduced exposure can shield from antibody binding, and provides more time for the virus to bind to ACE-2 before getting neutralized. The neutralization of WT-Omi S2 by S309 and 4A8 was substantially increased when antibody and virus was incubated for longer period of time whereas such increase in neutralization was much smaller for WT. The observation was further bolstered by reverting S2 mutations in Omicron spike which led to increased neutralization by RBD-antibodies. This provides an explanation why Omicron and all its subvariants from BA.1 to XBB retained S2 mutations while changing the mutational landscape in S1.

Omicron not only adapted to resist host NAbs but also varies from earlier VOCs with respect to TMPRSS2 usage and the entry mechanism. In contrast to previous VOCs, Omicron takes the endosomal route for entry. The mutations in S2 near the second cleavage site could affect its cleavage by TMPRSS2 and thereby fusion at the PM. Omicron entry depended on intracellular cathepsins but not TMPRSS2 whereas WT used TMPRSS2 and fused efficiently at PM(49, 62). This difference is substantially accounted for by the spike S2 mutations, particularly N856K and L981F substantially reduced the fusogenicity and attenuated the entry efficiency of Omicron. These two mutations are reverted to WT in Omicron subvariants (BA.2 to XBB) probably to regain the entry efficiency. Indeed, the fusogenicity and replication rate is enhanced from BA.1 to BA.5 with optimization of mutation profile (epistatic control) to adopt beneficial traits (63). The compensatory epistasis is more prevalent for maintaining ACE-2 affinity brought about by mutations within RBD(45). Recently, Kimura et al. showed the importance of S375F substitution for neutralization resistance to vaccine sera which also partially reduced Omicron’s fusogenicity(64). It is plausible that mutations in Omicron were selected for resistance to host NAbs by directly and indirectly modulating the epitopes in spike protein. As a result, multiple mutations stabilize spike trimer, retain affinity with ACE-2, but also alter the route of virus entry as an unintended consequence. These properties confer advantage in terms of better transmissibility in the face of host defense and might expand the tissue tropism (65). In the continued course of evolution, the genetic drift shapes Omicron spike further to be more resistant and fusogenic as exemplified by sub-lineages BA.2, BA.4, and BA.5 (50). The N856K and L981F in WT reduced fusogenic capacity of spike and increased E64D sensitivity – the propensity for endosomal entry. The lack of these two mutations in BA.2 sub-variant of omicron may explain why it is more fusogenic than BA.1 (50). Thus, present work delineated the role of mutations in spike-S2 subunit in the long-range interactions controlling the exposure of RBD and NTD and provide insights in conformational dynamics and allosteric modulation.

## Materials and methods

### Cells

Human embryonic kidney (HEK293T) cells were used for the transient expression of SARS-CoV-2 spike and HIV-1 proteins to prepare pseudotyped virus. HEK-293T cells expressing human angiotensin converting enzyme-2 (hACE2) or expressing hACE2 and TMPRSS2 were obtained from BEI Resources and used for the neutralization assays and infectivity assays. Vero-E6-TMPRSS2 (JCRB cell bank, JCRB #1818) and 293T-hACE2-TMPRSS2 cells were used in the cell-to-cell fusion assay. All cell lines were maintained at 37°C and 5% CO_2_ in DMEM supplemented with 10% fetal bovine serum (FBS).

### Serum samples and ethical approval

Serum samples were collected from individuals who were given two doses of Covishield or Covaxin vaccine. Sera were also collected from covaxin or covishield vaccinated individuals after the breakthrough infections in the third wave in February 2022 in India. Written informed consent forms were collected from all participants before collecting blood samples for sera. The study was approved by Institutional Ethics Committee of CSIR-Institute of Microbial Technology (IEC August 2020#2) and CSIR-Institute of Genomics and Integrative Biology (Approval no. CSIR-IGIB/IHEC/2020−21/01).

### Spike gene constructs

The SARS-CoV-2 spike constructs expressing the spike protein of Wuhan strain with D614G, Omicron (B.1.1.529), and Omicron BA.5 spike were based on the codon-optimised spike sequence of SARS-CoV-2 and were generated by GenScript Inc. Wuhan strain sequence with D614G was termed as wild type (WT) in this paper. All the spike constructs had 19 amino acid deletion at the C-terminal end for better cell-surface expression(66). The chimeric spike constructs were made between WT and Omicron genes where S1 or S2 of wild type spike were replaced by corresponding fragments of Omicron. S1 or S2 of Omicron fragments were PCR amplified by using specific primers. Similarly, vector backbone of WT spike excluding S1 or S2 was amplified by reverse primers with the overlap of 18 bases between insert-amplifying and vector-backbone-amplifying primers. The PCR fragments were digested by DpnI to remove template DNA and purified by gel-extraction. The vector and insert PCR fragments were then ligated by using In-Fusion cloning kit (CloneTech, Inc.). Two Omicron mutations N679K, P681H adjacent to furin cleavage site (FCS) were added in WT spike by using specific primers containing that sequence change and site-directed mutagenesis was done by overlapping PCR and ligation of insert and vector backbone was done by using In-Fusion cloning kit.

### Antibodies

The codon-optimised DNA sequences of heavy and light chain of S309, 4A8, CC40.8, and CV3.25 were purchased from GenScript and cloned in pcDNA3.4. The expression plasmids of COVA2-15, COVA2-17 were gifted by Rogier Sanders and Marit van Gils and described in Brouwer et al (67). The expi293 cells were co-transfected by Heavy and Light chain plasmids in 1:1 ratio by using polyethylenimine (PEI) and supernatants were harvested after 4 days post-transfection. The antibodies were purified from supernatants by using Protein A/G beads and finally stored in PBS for use in experiments.

### SARS-CoV-2 pseudovirus preparation and Neutralization Assay

Pseudovirus neutralization assays were performed by using HIV-1-based pseudovirus as described previously with some modifications (54). Briefly, HEK293T cells were transiently transfected with plasmid DNA pHIV-1 NL4·3Δenv-Luc and Spike-Δ19-D614G by using profection mammalian transfection kit (Promega Inc.). After 48 hours, culture supernatant was harvested and filtered using 0.22 μm filter, and stored at −80 °C until further use. 293T-hACE-2 (BEI resources, NIH, Catalog No. NR-52511) cells expressing the ACE2 receptors were cultured in DMEM (Gibco) supplemented with 5% FBS (Fetal Bovine Serum), penicillin−streptomycin (100 U/mL). Vaccine sera were heat inactivated at 56°C for 30 minutes and serially diluted in growth medium starting from 1:20 for neutralization of pseudovirus infection. The pseudovirus (PV) was incubated with serially diluted sera in a total volume of 100 μL for 1 h at 37°C. The cells were then trypsinized and 1 × 10^4^ cells/well were added to make up the final volume of 200μL/well. The plates were further incubated for 48 h in humidified incubator at 37 °C with 5% CO_2_. After incubation, neutralization was measured as indicator of luciferase activity in the cells (relative luminescence unit) by using nano-Glo luciferase substrate (Promega Inc.). Luminescence was measured by using Cytation-5 multimode reader (BioTech Inc.). The luciferase activity, measured as relative luminescence units (RLU), in the absence of sera was used as 100% infection. The serum dilution that resulted in half-maximal neutralization of pseudovirus (ID50) relative to no-serum control was determined from neutralization curves. All the assays involving the use of SARS-CoV-2 pseudoviruses were conducted after the approval by Institutional Biosafety Committee (CSIR/IMTECH/IBSC/2020/J21).

### Fusogenicity of spike protein

HEK293T cells were seeded in 6-well plate each well containing 4 x 10^5^ cells. Next day, cells reached ∼40-50% confluency and were co-transfected by using spike-expressing and GFP-expressing constructs. On the same day, in separate 96-well plate Vero-TMPRSS2 cells were seeded each well receiving 2×10^4^ cells. After 24 hours post-transfection, 293T cells were checked under a fluorescent microscope for qualitative assessment to confirm if the transfection of each spike construct was equivalent across various spike constructs. The cells were then trypsinised and 5 x 10^3^ cells/well were added on the monolayer of Vero-TMPRSS2 cells in 96-well plate. The plate was incubated at 37°C and periodically checked under microscope for syncytia formation. The size of the syncytia was calculated using NIS Elements software (Nikon Instruments Inc.)

### Infectivity Assay

1 x 10^4^ 293T-hACE2-TMPRSS2 cells/well were seeded in 96-well plate. Next day, pseudoviruses stored at -80°C were thawed at room temperature. The serial dilution of virus was done in growth medium starting from neat up to 128-fold dilution. The growth medium from 293T-hACE2-TMPRSS2 cells were removed and 100µL virus was slowly added. The plate was incubated for 48 hours at 37°C in a CO_2_ incubator and luciferase activity was quantified by adding nano-glo luciferase substrate (Promega). The luminescence was recorded by using Cytation 5 plate reader (BioTek inc). Based on the titration curve, the dilution at which 2 x 10^6^ RLU was obtained was identified for each variant and was considered to contain equivalent infectious units. With this normalization, the infectivity of pseudovirus was measured in serial dilution of pseudovirus in 293T-hACE2-TMPRSS2 cells.

### Infectivity in the presence of protease inhibitors

The sensitivity to protease inhibitors - camostat or E64d – was assessed in 293T-ACE2-TMPRSS2 cells. 2×10^4^ cells/well were plated in 96-well plate. Next day, protease inhibitors were serially diluted starting from 40µM in 100ul growth media. The media on the cells was removed and the media containing inhibitors was added on the cells and incubated for 2 hours. 100ul virus stock was then added on top to make the final volume 200ul and plate was further incubated for 48 hours. The infection was measured by assaying activity of intracellular luciferase enzyme by using nano-go luciferase system (Promega).

### RBD-based immunogen and mice sera

All mice sera raised against WT RBD sequence were used from our previously published study (53, 54). The immunogens were based on WT-RBD or Omicron-RBD and were expressed in expi293 cells and purified by using Ni-NTA column followed by tag removal by digesting with protease. Animals were immunized with 10 ug of RBD-based subunit vaccine candidates mixed with adjuvants in 50ul injection volume intramuscularly, with the prime dose at day 0 and a boost on day 21. After two weeks of complete immunization blood was collected and serum was separated and stored at -80°C until further use. All animal studies were approved by the Institutional Animal Ethics committee (IAEC) of Indian Institute of Science.

### FACS staining of spike proteins expressed on the transfected cells

HEK293T cells (0.6 x 10^6^ cells/well in 6-well plate) were transfected with spike-expressing construct. After 24 hours, cells were trypsinised and washed with PBS and subsequently resuspended in FACS buffer (PBS containing 5% FBS). The cells were stained with spike-specific monoclonal antibodies (COVA 2-15, S309, and 4A8) 4-fold serially diluted in FACS buffer starting from 10ug/mL. The cells were incubated with antibodies for 1 hour at 4°C. The cells were washed twice with FACS buffer followed by incubation with 100uL goat anti-Human IgG, Alexa Fluor^TM^ 488, at 1:500 dilution of original stock for 45 minutes at 4°C. The cells were washed twice, resuspended in FACS buffer, and the fluorescence intensity was measured using BD FACSVerse^TM^. The data was analysed in FlowJo software (Treestar).

**Figure S1:**
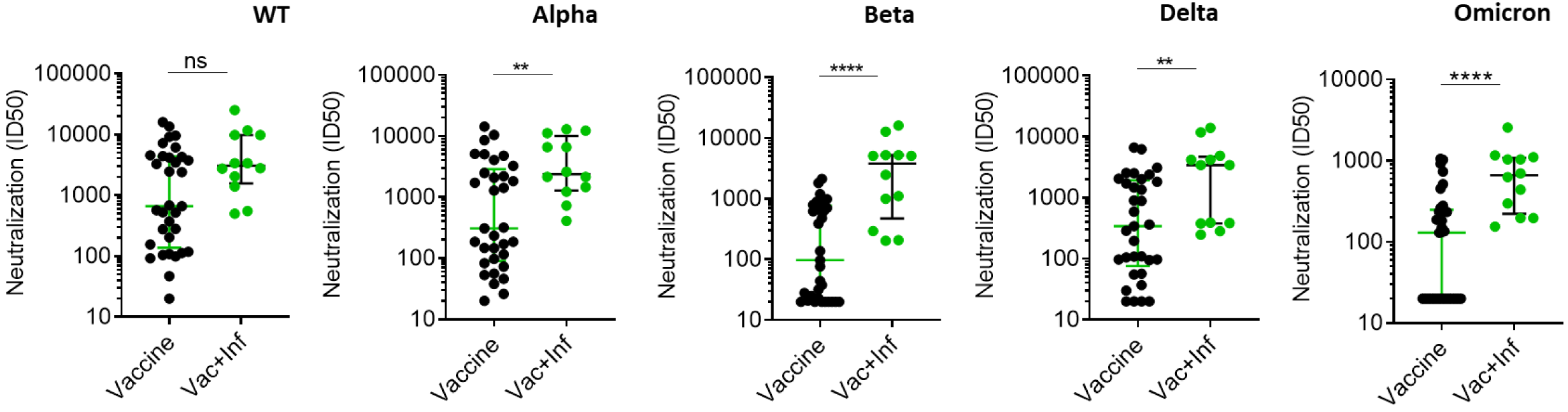
Nuetralization sensitivity of SARS-CoV-2 pseudoviruses to vaccine and vaccine plus infection sera: Shown are the neutralization titers for various pseudoviruses against the sera obtained after vaccination (black circles) and from the individuals who got infection in the third wave after vaccination (green circles). Statistical significance is indicated by using Mann Whitney test.

**Figure S2:**
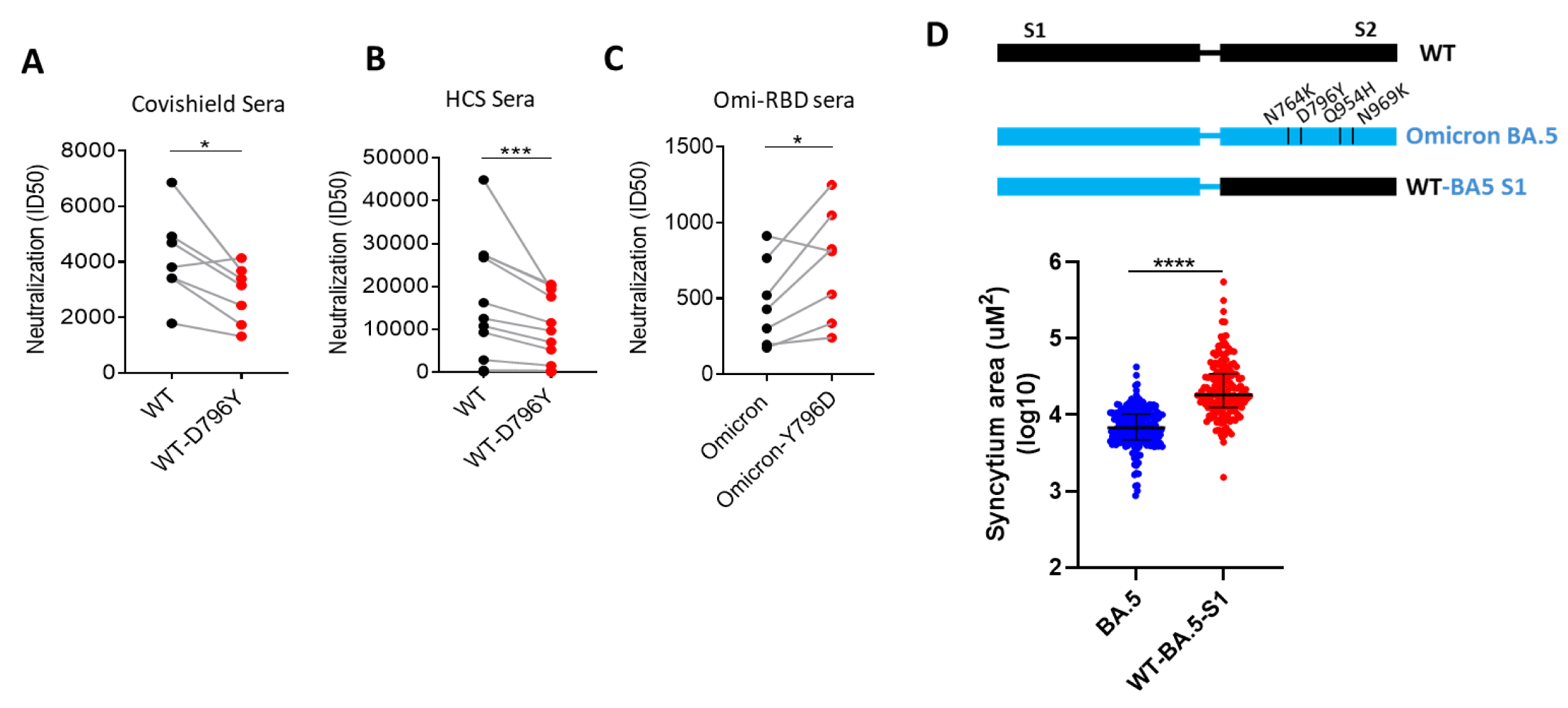
Modulation of neutralization sensitivity by D796Y mutation: Neutralization titers against WT and WT-D796Y for A) Covishield vaccine sera B) human convalescent sera. C. Neutralization sensitivity of Omicron BA.1 and Omicron-Y796D to mouse sera raised against Omicron-RBD. D. The fusogenicity of Omicron BA.5 and WT-BA5 S1 and statistical significance shown using Mann-Whytney test.

**Figure S3:**
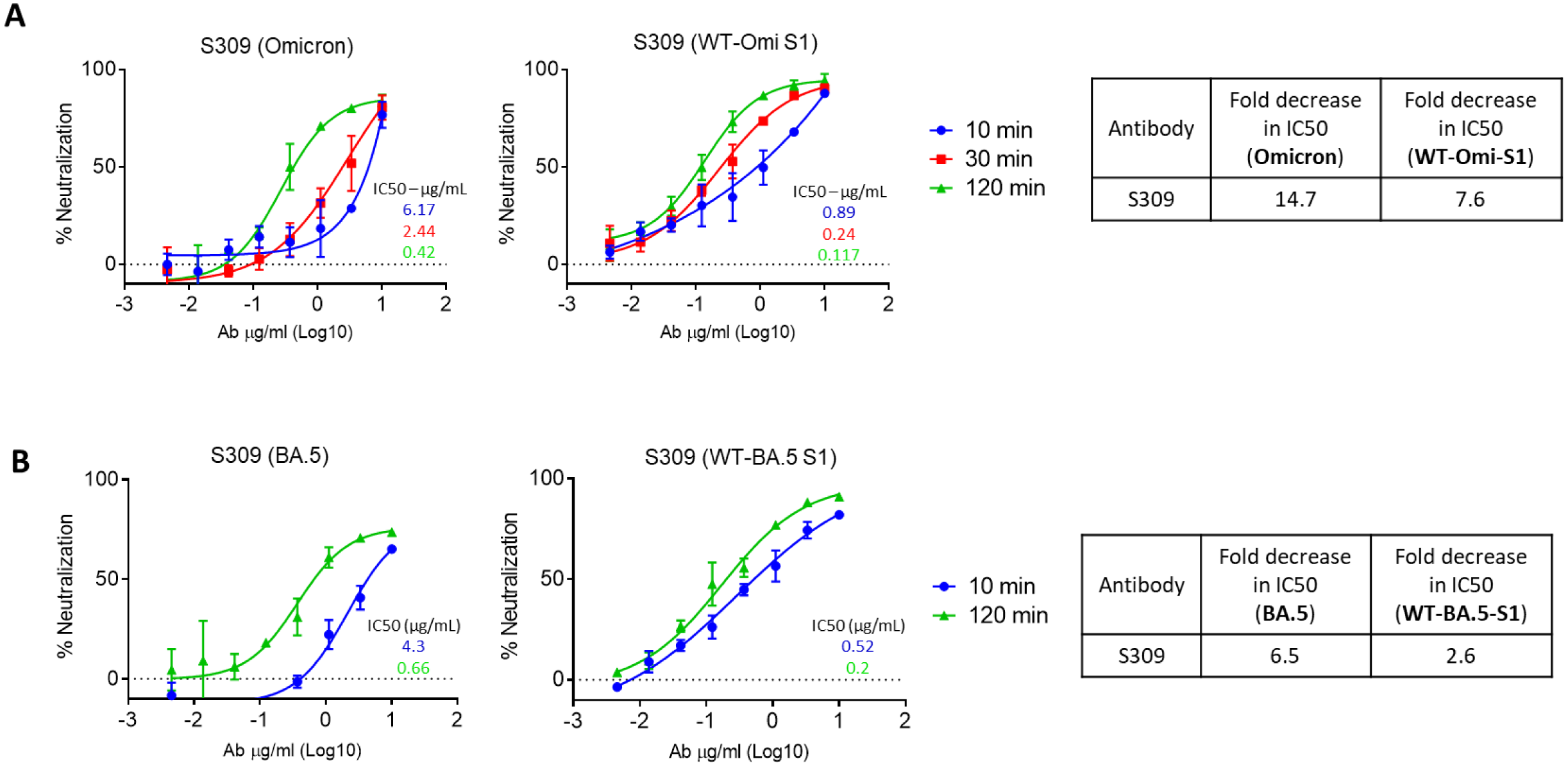
Time-dependent neutralization of pseudoviruses by S309: A. Neutralization of the Omicron and WT-Omi S1 pseudoviruses at incubation time of 10 or 30 or 120 minutes. The IC50 values (ug/ml) are shown for neutralization curves and indicated by same color for respective neutralization curves. The fold decrease in IC50 value from incubation period 10 min to 120 minutes is given in the table. B. Neutralization of BA.5 and WT-BA.5 S1 for 10- or 120-minute incubation period. The fold decrease in IC50 as in A is given in the corresponding table.

**Figure S4:**
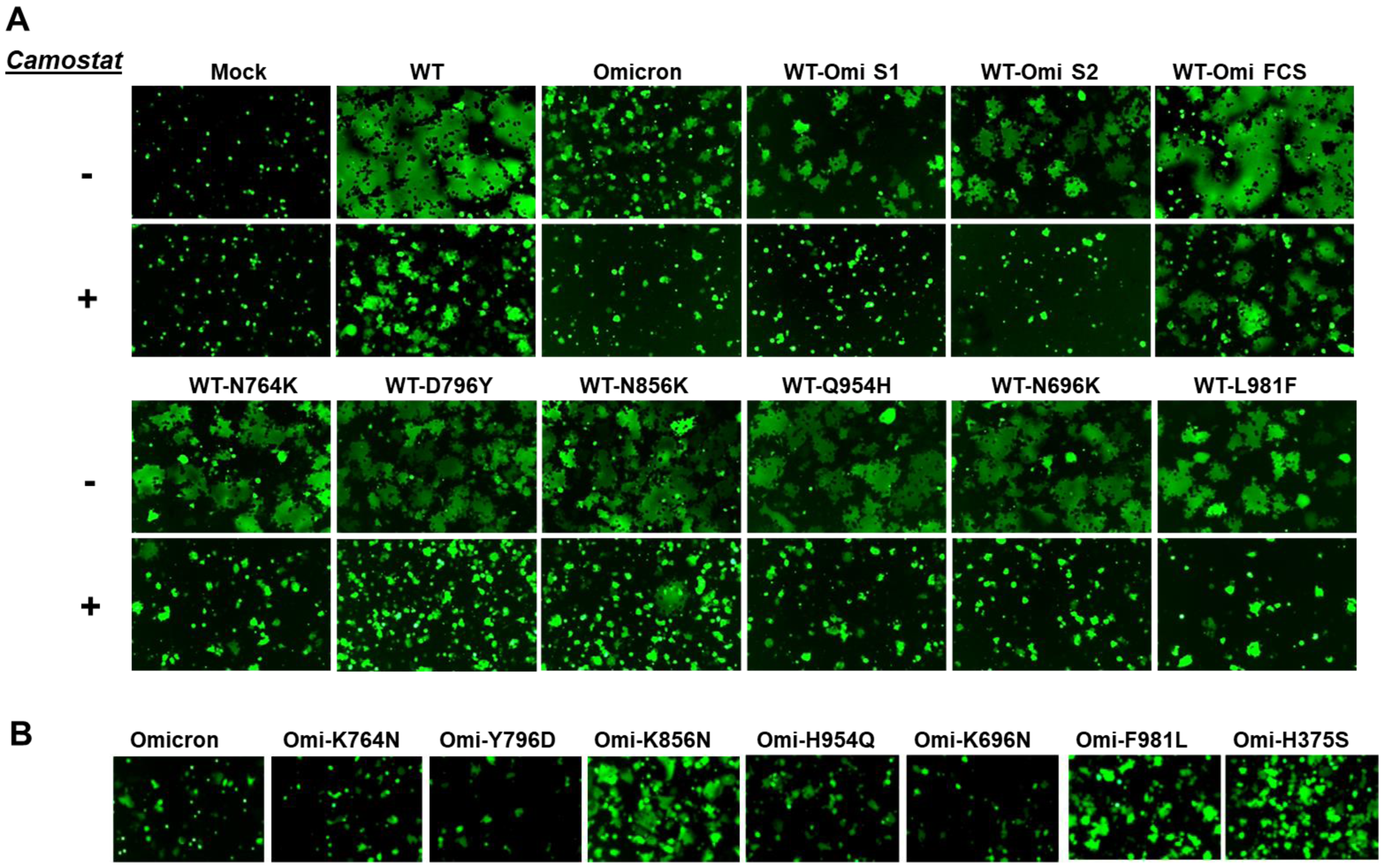
Fusogenicity of Spike variants in veroE6-TMPRSS2 cells: A. Shown are the images of syncytia formed after addition of transfected 293T cells on the monolayer of VeroE6-TMPRSS2 cells which were either pre-treated or not with camostat. The images were taken after 3 hours of addition of cells. Mock represents the cells transfected only with GFP-expressing construct but not Spike. B. The cell-cell fusion of Omicron and its mutants post 3-hour incubation

## Author Contributions

RPR conceived the study. RPR, SK, RD, DC, KK, RS, NS performed research. RD, DC performed molecular cloning. RPR, SK, RD, NS analysed the data. RV designed RBD-based immunogens for the immunogenicity in mice and supervised the immunogenicity studies. RD and RS expressed and purified Spike and RBD proteins and antibodies. RS performed the mouse immunizations and serum collection. S.S, A.T., S.J., R.P. provided sera from COVID-19 human convalescent sera from the first wave of pandemic, and vaccinated individuals. RPR wrote and RV edited the paper.

## Conflict of Interest

RV and RS are inventors on a patent application for RBD antigens used in this study, other authors declare no conflict of interest.

## Acknowledgements

We thank Dr. Paul Bieniasz for providing SARS-CoV-2 Spike expressing plasmid and pHIV-1-NL4.3-nanoLuc construct. Dr. Rogier Sanders and Dr. Marit van Gils for providing H and L chain constructs of SARS-CoV-2 neutralizing antibodies COVA2-15 and COVA2-17. We thank BEI Resources for providing HEK293T-ACE2 and HEK293T-ACE2-TMPRSS2 cells. This work was funded by a grant from Science and Engineering Research Board (IPA/2020/000168 to RPR) and Council of Scientific and Industrial Research (CSIR) to KGT and RPR, and Bill and Melinda Gates Foundation (INV-042471) to R.V. RV also acknowledges infrastructural support from the following programs of the Government of India: DST-FIST, UGC Center for Advanced Study, MHRD-FAST, the DBT-IISc Partnership Program, and of a JC Bose Fellowship from DST. DC and KK thank CSIR and IISc respectively for doctoral fellowship. RPR is the recipient of DBT-Ramalingaswami fellowship. R.P. acknowledges the support of CSIR-IGIB grant (MLP-2005) and Fondation Botnar (CLP-0031).

